# *S*-nitrosylation affects TRAP1 structure and ATPase activity and modulates cell response to apoptotic stimuli

**DOI:** 10.1101/859629

**Authors:** Fiorella Faienza, Matteo Lambrughi, Salvatore Rizza, Chiara Pecorari, Paola Giglio, Juan Salamanca Viloria, Maria Francesca Allega, Giovanni Chiappetta, Joëlle Vinh, Francesca Pacello, Andrea Battistoni, Andrea Rasola, Elena Papaleo, Giuseppe Filomeni

**Author notes:** Facultat de Farmàcia, Universitat de Barcelona, 08028, Barcelona, Spain (Juan Salamanca Viloria). Cancer Research UK Beatson Institute, Garscube Estate, Switchback Road, Glasgow G611BD, UK. To whom correspondence should be addressed: Elena Papaleo, Giuseppe Filomeni,;.

## Abstract

The mitochondrial chaperone TRAP1 has been involved in several mitochondrial functions, and modulation of its expression/activity has been suggested to play a role in the metabolic reprogramming distinctive of cancer cells. TRAP1 posttranslational modifications, i.e. phosphorylation, can modify its capability to bind to different client proteins and modulate its oncogenic activity. Recently, it has been also demonstrated that TRAP1 is *S*-nitrosylated at Cys501, a redox modification associated with its degradation *via* the proteasome. Here we report molecular dynamics simulations of TRAP1, together with analysis of long-range structural communication, providing a model according to which Cys501 *S*-nitrosylation induces conformational changes to distal sites in the structure of the protein. The modification is also predicted to alter open and closing motions for the chaperone function. By means of colorimetric assays and site directed mutagenesis aimed at generating C501S variant, we also experimentally confirmed that selective *S*-nitrosylation of Cys501 decreases ATPase activity of recombinant TRAP1. Coherently, C501S mutant was more active and conferred protection to cell death induced by staurosporine. Overall, our results provide the first *in silico, in vitro* and cellular evidence of the relevance of Cys501 *S*-nitrosylation in TRAP1 biology.

## 1. INTRODUCTION

Tumor necrosis factor type 1 receptor-associated protein 1 (TRAP1) is a molecular chaperone belonging to the Hsp90 family, which shares 34% identity and 60% similarity with the other orthologs [1]. Unique in the Hsp90 family, TRAP1 shows a predominant, but not exclusive mitochondrial localization [2]. Indeed, it is the only member showing a 59 aa mitochondrial import sequence, which is cleaved in the organelle [3].

TRAP1 acts as a homodimer, although it has recently been proposed it is mostly present under tetrameric form, with this quaternary structure modulating mitochondrial oxidative phosphorylation [4]. Each subunit presents a N-terminal domain (NTD) domain, involved in ATP binding and hydrolysis; a middle-domain (MD), that contains that contains part of the ATP-binding pocket [5], and the binding site for client proteins; a C-terminal domain (CTD) that constitutes the interface for homodimerization [6]. Differently form the other Hsp90 orthologs, TRAP1 lacks the linker domain between MD and CTD, and exhibits a N-terminal extension that acts as thermal regulator of its chaperone activity [7].

The ATPase cycle of TRAP1 has been characterized in detail, with both protomers assisting the folding of the target proteins through structural modifications associated with repeated cycles of ATP binding, hydrolysis and release [2,8,9]. During the ATPase cycle, TRAP1 can be present under three different states: *i*) an open conformation (called *apo*); *ii*) a close conformation with the NTD placed between the two protomers; *iii*) an intermediate coiled-coil conformation with both NTD in close proximity. In the absence of substrates, TRAP1 is present in an open state, while the binding of two 2 molecules of ATP induces structural changes leading to a closed asymmetric conformation [8], this being distinctive of TRAP1 among Hsp90 family members. ATP binding provides the energy required for substrate remodelling which takes place by a two-step reaction. The hydrolysis of the first ATP produces symmetric changes in protomers, establishing a structural rearrangement in the substrate binding site. The hydrolysis of the second ATP leads to an ADP-bound state. This conformation is associated with a compact structure of the chaperon and is preparatory for the release of the remodeled substrate along with ADP [5,6,8,10].

Mitochondrial TRAP1 expression differs among tissues (e.g. brain, skeletal muscle, liver, kidney, heart and pancreas) [1] and – with the exception of the brain – it is kept low [11], whereas it is found abnormally increased in tumors [12,13]. In the last few years it has become clear that, besides mitochondrial proteostasis [14], TRAP1 plays key roles also in several cellular processes, such as cell death regulation [15], antioxidant response [16]; mitochondrial transmembrane potential control [12, 13] and metabolism tuning [17]. In regards to the last process, it has been demonstrated that TRAP1 regulates the metabolic rewiring from oxidative phosphorylation to aerobic glycolysis (the so-called Warburg effect) through its inhibitory activity on complex IV and, mostly, on complex II of the electron transport chain [12], this finally leading to the stabilization of hypoxia-inducible factor 1α (HIF1α) [18].

TRAP1 activity is regulated by posttranslational modifications. In particular, phosphorylation events mediated by PTEN induced kinase 1 (PINK1) have been reported to be instrumental for antioxidant and anti-apoptotic response [19], whereas those induced by the extracellular signal-regulated kinase 1 and 2 (ERK1/2) regulate TRAP1 ability to modulate cancer metabolism [20]. Similarly, SIRT3-mediated deacetylation has been very recently proposed to have a role in rewiring glioblastoma stem cell metabolism [21]. In addition to phosphorylation and (de)acetylation, it has been also demonstrated that TRAP1 undergoes *S*-nitrosylation at Cys501, which is preparatory for its degradation *via* the proteasome [22]. C501S mutant of TRAP1 is, indeed, more stable in conditions of excessive *S*-nitrosylation, e.g., in cells lacking the denitrosylase *S*-nitrosoglutathione reductase (GSNOR).

Here we predict using biomolecular simulations that *S*-nitrosylation modifies the functional motions of TRAP1, as well as its intra- and intermolecular interaction network and activity. We also observe that TRAP1 *S*-nitrosylation at Cys501 affects mitochondrial homeostasis and modulates response to staurosporine-induced cell death, indicating a possible biological role for this modification.

## 2. MATERIALS AND METHODS

The scripts, inputs and output files for the computational parts of the work are freely available in a GitHub repository associated with our publication (https://github.com/ELELAB/TRAP1_activity) and the trajectories have been deposited in OSF (https://osf.io/wytg2).

### 2.1. Molecular dynamics simulations

We used the full-length *Danio rerio* TRAP1 (PDB entry: 4IPE, resolution 2.29 Å, [5]) as a starting structure for the molecular dynamics (MD) simulations, following the preparation of the structure used in a recent work by Colombo’s group [23]. We selected this structure since it is the only full-length structure available for TRAP1 and the one used in structural studies carried out so far. Moreover, we used MODELLER 9.15 [24] to generate a model of each individual protomer of TRAP1 using as template the full-length *D. rerio* structure (chain A and B from 4IPE) in which we modeled the missing residues T153, D240-A241, V370-G375, T567-E587, R617, T639-I652, K718-H719 in chain A and A149-D152, A201-A208, E273-S376, T389-T392, Q640-Q651, K718-H719 in chain B. We removed the AMP-PNP molecule from the crystallographic structure before carrying out the simulations to keep our model system the simplest possible. This strategy is also based on the notion that functional dynamics patterns are intrinsic to protein structures [25]. Cofactors or binding of other biomolecules generally promote population shift of already existing conformational states of a protein ensemble [26]. The orthologous protein in *D. rerio* has an 80% sequence identity when compared to the human TRAP1 variant and no structures are available for the full-length human variant to the best of our knowledge.

Moreover, due to the known TRAP1 structural asymmetry [5,10], we performed simulations of both the dimeric assembly and of each individual protomer (chain A and B from 4IPE, respectively). For each individual protomer, we removed their first 23 residues (T85-A107) corresponding to the N-terminal strap that wraps around the other protomer in the dimeric assembly to avoid that the floppy terminal disordered regions engaged in non-native interactions with the rest of the protomer. Chain A and B correspond to the buckled and straight protomers, respectively, as defined in [10]. For each protomer, we carried out MD simulations of the unmodified (TRAP1_SH_) and *S*-nitrosylated variant on cysteine 516 (TRAP1_SNO_). The SNO variant was obtained by Vienna-PTM [27]. Dimer simulations were performed only for the SH variant. The details of the simulations used in the study are reported in **Table 1**.

**Table 1.** Summary of the MD simulations collected in this study.

| VARIANT | STARTING STRUCTURE | FORCE FIELD | SIMULATION LENGTH (ns) | CYS 516 |
| --- | --- | --- | --- | --- |
| TRAP1<sub>SH</sub> | Chain A, 4IPE | ff99SB*-ILDN/TIP3P | 150 | reduced |
| TRAP1<sub>SH</sub> | Chain A, 4IPE | GROMOS 54a7/SPC | 150 | reduced |
| TRAP1<sub>SNO</sub> | Chain A, 4IPE | ff99SB*-ILDN/TIP3P | 150 | S-nitrosylated |
| TRAP1<sub>SNO</sub> | Chain A, 4IPE | GROMOS 54a7/SPC | 150 | S-nitrosylated |
| TRAP1<sub>SH</sub> | Chain B, 4IPE | ff99SB*-ILDN/TIP3P | 150 | reduced |
| TRAP1<sub>SH</sub> | Chain B, 4IPE | GROMOS 54a7/SPC | 150 | reduced |
| TRAP1<sub>SNO</sub> | Chain B, 4IPE | ff99SB*-ILDN/TIP3P | 150 | S-nitrosylated |
| TRAP1<sub>SNO</sub> | Chain B, 4IPE | GROMOS 54a7/SPC | 150 | S-nitrosylated |
| TRAP1<sub>SH</sub> | Dimer, 4IPE | ff99SB*-ILDN/TIP3P | 500 | reduced |
| TRAP1<sub>SH</sub> | Dimer, 4IPE | GROMOS 54a7/SPC | 500 | reduced |

We performed the simulations using GROMACS version 4.6 [28], employing the force fields ff99SB*-ILDN [29] and GROMOS 54a7 [30] for which parameters for *S-*nitrosylation were available [31,32]. The protein was solvated in a dodecahedral box of water molecules in periodic boundary conditions at a concentration of 150 mM NaCl, neutralizing the net charge of the system. We used Single Point Charge (SPC) or the TIP3P solvent models for GROMOS and AMBER force fields, respectively. We defined the protonation state of each histidine using ProPka and the PDB2PQR server (https://academic.oup.com/nar/article/35/suppl_2/W522/2920806) [33] and modeled all the histidine in the Nε tautomeric state with the exception of H326 and H573 in the Nδ state. The system was equilibrated through a series of energy minimization, solvent equilibration, thermalization and pressurization steps (see GitHub repository associated with the publication for details). We performed the productive MD simulations in the canonical ensemble at 298 K. We collected 150 and 500 ns for monomers and dimers, respectively. We used the Particle-mesh Ewald summation scheme and we truncated Van der Waals and short-term Coulomb interactions at 9 Å. For more details, we provided the parameter input files iin the GitHub repository associated with the publication the parameter input files.

### 2.2. Protein Structure Network

We used a contact-based protein structure network (PSN) to analyze the MD ensemble of the TRAP1_SH_ dimer, as implemented in PyInteraph [34]. For the contact-based PSN, we selected a 5 Å distance cutoff to define the existence of a link between the nodes, which is a suitable compromise between an entirely connected and a sparse network, according to our recent work on PSN cutoffs [35]. The distance was estimated between the side-chain center of masses (excluding glycines). Since MD force fields have different mass definitions, we used the PyInteraph mass databases compatible with the two force fields selected for this study. We retained in the PSN for analysis only those edges which were present in at least 20% of the simulation frames to remove spurious interactions, as previously applied to other case studies [34]. Prior to the analysis, we evaluated the time evolution of the root mean square deviation (rmsd) of the NTD, MD and CTD, respectively. We observe that approximately 120 and 50 ns were needed to have a stable rmsd profile over time (see GitHub repository for the rmsd analyses). We thus carried out the PSN analyses only on the portion of the MD trajectories with stable rmsd profiles.

We defined as hubs those residues of the network with at least three edges, as commonly done for networks of protein structures [36]. We applied a variant of the depth-first search algorithm to identify the shortest path of communication between the SNO site of each monomer (C516) and all the other residues of the protein (testing different cutoffs for the path lengths to be sure we cover also very distal effects, i.e. 15, 25 and 40). We focused on the analyses of the paths with a maximal path length of 25 since increasing this value to 40 did not provide additional target residues. We defined the shortest path as the path in which the two residues were non-covalently connected by the smallest number of intermediate nodes. We did not include in the path analyses, the residues in the regions 429-485 and 543-550 of the protomer where the source SNO site was located since they contained several residues within 8 Å of distance from C516 (i.e., not suitable as distal sites). All the PSN and path calculations have been carried out using the PyInteraph suite of tools [34].

### 2.3. FoldX calculations

We employed the FoldX energy function [37] to estimate the impact of mutation to serine at the C516 site on structural stability. The calculations resulted in an average ΔΔG (differences in ΔG between mutant and wild-type variant) for each mutation over five independent runs performed using the same structure used for MD simulations (PDB entry 4IPE), along with some of the available structures for the TRAP1 human variant (i.e. PDB codes here 4Z1F, 4Z1G, 4ZIH, and 5HPH). For the X-ray structures of TRAP1 dimers (4IPE and 5HPH), we estimated the ΔΔG values the isolated protomers, i.e. chain A and chain B. The typical prediction error of FoldX is about 0.8 kcal/mol [37]. Twice the prediction error (i.e., 1.6 kcal/mol) was used as a threshold to discriminate between neutral and deleterious mutations in the analyses.

### 2.4. Principal Component Analysis (PCA)

We employed a dimensionality reduction technique based on Principal Component Analysis (PCA) [38] to represent and compare in a conveniently simplified two-dimensional (2D) space the conformational space explored by the two different TRAP1 variants (TRAP1_SH_ and TRAP1_SNO_). PCA, applied to MD trajectories, allows to calculate the eigenvectors (principal components, PCs) of the mass-weighted covariance matrix of the atomic positional fluctuations. We calculated the covariance matrix of only Cα atoms of the protein for the region 108-716. We did not include the unstructured regions at the N- and C-terminal of the protein, whose pronounced motions might otherwise hide important differences in the rest of the protein. We aligned the structures from the concatenated trajectories, before the covariance matrix calculation, on a subset of Cα atoms that do not undergo pronounced conformational changes during the simulations. In details, we selected for the alignment the following residues: 591-608, 611-614, 618-626, 630-638, 656-659, 664-675, 677-695 in the CTD of the buckled protomer.

We used a common essential subspace derived from a PCA of the concatenated trajectories of TRAP1_SH_ and TRAP1_SNO_. In details, we used two concatenated trajectories, each of them including all the MD simulations of TRAP1_SH_ and TRAP1_SNO_ protomers and performed with one force field (ff99SB*-ILDN or GROMOS 54a7). The first two principal components explains more than 77.6 and 83.9 % of the total variance for ff99SB*-ILDN and GROMOS 54a7, respectively. We thus obtained a 2D density plots along the first two principal components, using an in-house R script that performs 2D kernel density estimation, employing the *kde2d* function from the *MASS* package and the *stat_density_2d* function from *ggplot2* package. We collected the conformations of TRAP1 in the concatenated trajectories corresponding to the most densely populated areas identified on the PCA 2D plots.

### 2.5. Plasmids

TRAP1 WT and C501S were cloned in pET-26b(+) plasmid for bacterial expression and in pcDNA3.1(+)-C-eGFP plasmid for mammalian expression using the Gene Synthesis & DNA Synthesis service from GeneScript Biotech.

### 2.6. Purification of recombinant proteins

TRAP1^WT^ and TRAP1^C501S^ recombinant proteins were produced in BL21(DE3) *E. coli* cells after 3 h-induction with 1 mM IPTG (Isopropil-β-D-tiogalattopiranoside, VWR). Protein purifications were performed using Ni-NTA resin (Quiagen) according to manufacturer’s instructions. Proteins release from the resin packed in a FPLC column was achieved using a linear imidazole gradient generated by mixing buffer A (50 mM Phosphate Buffer, 250 mM NaCl, 10 mM imidazole, 10 mM β-mercaptoethanol, pH 7.8) and B (buffer A with 250 mM imidazole). Eluted fractions containing recombinant proteins were then collected and combined. After purification the proteins were dialyzed in the *storage buffer* (50 mM Tris-HCl, 150 mM NaCl, 1 mm DTT, 1 mM EDTA, pH 7.5) and stored into aliquots.

### 2.7. *S*-nitrosylation in vitro

Purified WT and C501S forms of TRAP1 (TRAP1^WT^ and TRAP1^C501S^) were dialyzed in *storage* buffer. Before treatments, DTT was removed by Zeba Spin Desalting Columns (Thermo Fisher) and proteins transferred in *reaction* buffer (NaCl 150 mM, TRIS 50 mM, pH 7.5). *S*-nitrosylation and denitrosylation were performed at 37°C in the dark by incubations with *S*-nitroso-*N*-acetilpenicilamine (SNAP - Enzo Life Sciences, Inc) 500 μM for 4 h and with ascorbate (Sigma-Aldrich) 20 mM for 30 min, respectively.

### 2.8. TRAP1 ATPase assay

TRAP1 ATPase activity was measured by by quantifying the release of inorganic phosphate as PO_4_ nmoles/TRAP1 nmoles/min. The ATPase assay was performed at 37°C for 1 h, incubating 5, 10, and 15 μg of proteins with 200 μM ATP and 10 mM MgCl_2_, in *reaction* buffer, in the dark. The amount of inorganic phosphate (PO_4_^3-^) released due to the activity of TRAP1 was measured using the Malachite Green Phosphate Assay Kit (Sigma-Aldrich), by detecting the increase in absorbance at 650 nm with a plate reader (Tecan).

### 2.9. Cell culture, treatments and transfections

HeLa cell line was obtained from the American Type Culture Collection (ATCC) and grown in DMEM (Thermo-Fisher Scientific) supplemented with 10% FBS (Thermo-Fisher Scientific) and 1% Penicillin/Streptomycin mix (EuroClone). Cells were maintained at 37°C in an atmosphere of 5% CO_2_.

Cells were transfected using polyethylenimine (PEI) (Sigma-Aldrich) in Opti-MEM medium (Thermo-Fisher Scientific), and incubated twice with plasmids and PEI with a 8 h-release between one transfection and the other. After the second transfection cells were incubated in DMEM and treated with 250 μM of the NO donor DETA-NONOate (Enzo Life Sciences) for 22-24 h and/or 200 nM Staurosporine (STS) (Sigma-Aldrich) for 4h.

### 2.10. Analysis of cell viability and cell death

Cell death was analyzed with the direct count of dead cells upon Trypan Blue staining (Sigma-Aldrich).

### 2.11. Analyses of mitochondrial transmembrane potential (Δψ_m_)

Mitochondrial Δψ_m_ was analyzed by incubating the cells with 200 nM tetramethylrhodamine methyl ester (TMRM, Thermo-Fisher Scientific) for 30 min in serum-free DMEM. Stained cells were washed twice with cold PBS, collected and analyzed by flow cytometry (FACs Verse, BD-biosciences).

### 2.12. Western blot

Samples preparation and acquisition of Western blots were performed as previously reported [22]. Protein concentration was determined by the method of Lowry [39]. Western blots shown are representative of at least n=3 independent experiments giving similar results. We used the following primary antibodies: anti-HSP75 (TRAP1) and anti-Actin from Santa Cruz Biotechnology; anti-Vinculin from Sigma-Aldrich; anti-PARP1 from Enzo Life Sciences; anti-Caspase 3 from Cell Signaling.

### 2.13. Statistical analyses

Values are expressed as means ± SEM and statistical significance was assessed by Student’s t-test or one-way ANOVA test using Prism 8.0 (GraphPad Software, Inc.) in order to determine which groups were significantly different from the others.

## 3. RESULTS AND DISCUSSION

### 3.1 The S-nitrosylation site establishes a long-range communication with the active site

As above introduced, the chaperone TRAP1 exploits its function as a homodimer, with each monomer consisting of three distinct domains: a N-terminal domain (NTD, residues 101-308) that harbors the ATP binding site, a middle domain (MD, residues 311-571) where the binding site for client proteins resides, and a C-terminal domain (CTD, 587-719) important for dimerization (**Figure 1A**).

**Figure 1.**
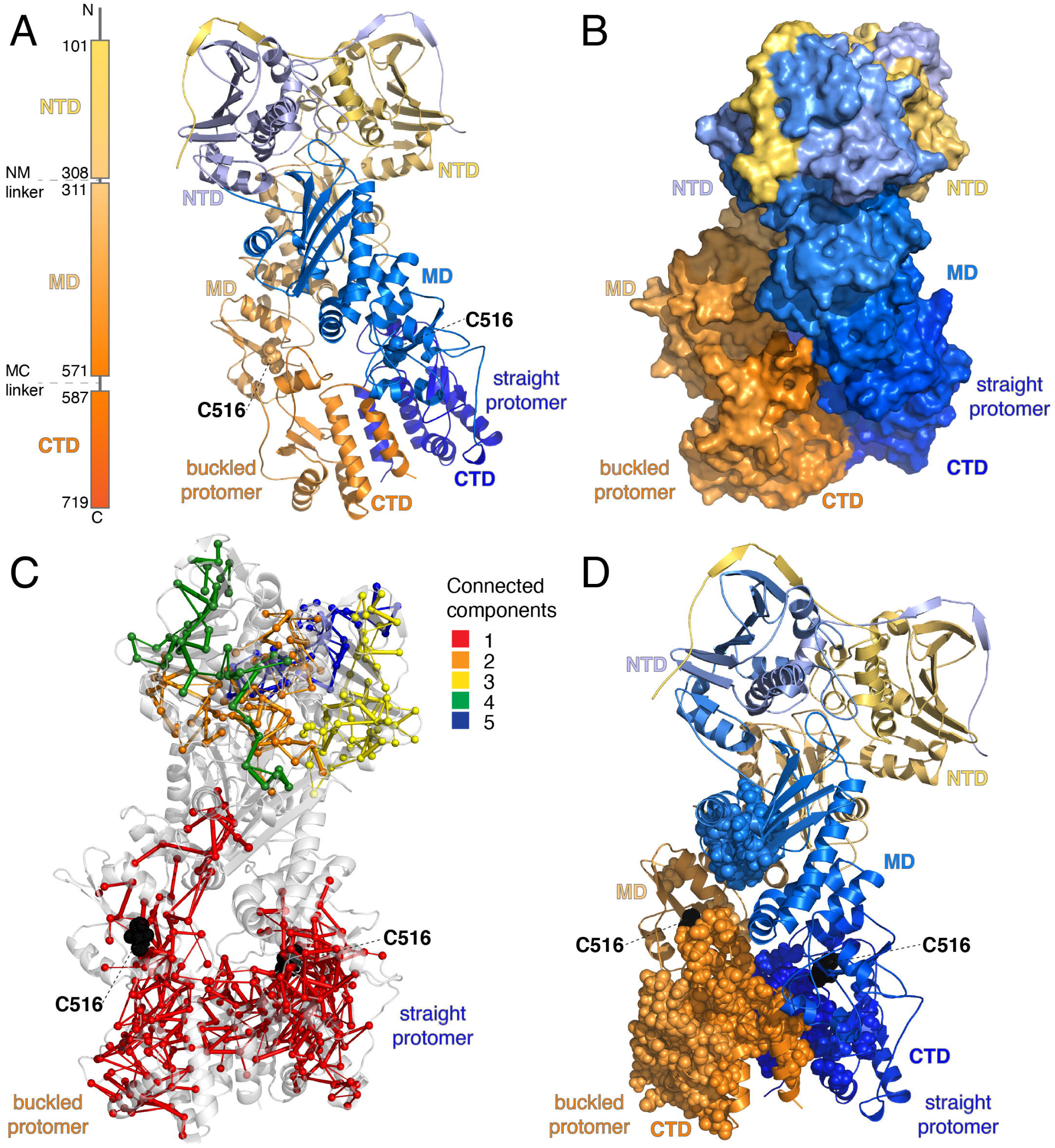
TRAP1 *S*-nitrosylation site modulates propensity for intra- and intermolecular communications at distal sites of TRAP1. **A)** The localization of C516 (*D. rerio* numbering) in the 3D structure of TRAP1 is shown with reference to the different domains. The two protomers are colored in different shades of yellow/orange (chain A from the PDB file 4IPE) and blue (chain B) to highlight the three structural domains. C516 is located in the middle domain. **B)** The buckled (orange) and straight (blue) protomers are shown as a reference. They correspond to chain A and B of the PDB file 4IPE, respectively. **C)** Contact-based protein structure network from the MD simulations of TRAP1_SH_ dimer. The results for the simulation with ff99SB*-ILDN force field are illustrated as an example. The data for the other simulation are reported in the GitHub repository associated with the publication. The C516 residues from both the protomers belong to the same and largest connected component of the network which embraces the regions at the interface between the protomers in MD and CTD, along with a part of the NTD of the straight protomer. **D)** Example for the path analysis using C516 of the buckled protomer as a source residue. MD simulation with the ff99SB*-ILDN force field is illustrated. The results for the other cases of study are reported in the GitHub repository associated with the publication. C516 features the propensity for long-range communication to different sites of both protomers, including the regions at the interface between MD and CTD where the client proteins bind or that are important hinges for the conformational changes of the chaperone. Moreover, the structural communication can also extend to the NTD of the cognate straight protomer.

The available crystal structures of TRAP1 and biophysical experiments carried out in *D. rerio* variant [5,10] revealed asymmetry between the two protomers with differences especially at the interface between MD and CTD, where the client protein binds. The two protomers have been defined as buckled and straight [5,10], which are illustrated in **Figure 1B** for sake of clarity. The SNO site is located at approximately 60 Å away from the active site in the middle domain of TRAP1 (**Fig. 1A**).

We thus employed a method to predict structural perturbations that can be propagated over long distances from one site to other sites of a protein, based on network theory [34,35]. The approach is based on the collection of an ensemble of conformations for the protein of interest using MD simulations, followed by the analysis of the conformational ensemble with a contact-based protein structure network (PSN). On the resulting network, an algorithm to search for the shortest path of communications between the site of interest is applied, according to the knowledge that in protein structures communication of structural effects among distal sites is preferred along the shortest roads [35]. In this work, we collected the MD simulations with two different force fields to assess the robustness of the results, namely the physical models used to describe the protein and its environment during the simulation. We used the *D. rerio* structure of TRAP1, as detailed in the method, where the *S*-nitrosylation site is the C516 (conserved in the human variant at position 501, C501). We noticed that the SNO site was in a region surrounded by hubs residues (see GitHub repository associated with the publication), which are important residues to organize the global network structure and transmit information from one site to the other. The SNO site is also included in the largest connected component of the PSN in both the simulations of the TRAP1_SH_ dimer (**Fig. 1C** in red). We calculated all the paths of long-range communication from each SNO site to the rest of the protein. We observe that each SNO site has the potential to exert distal effects within the corresponding protomer (i.e., C516 of the buckled protomer to distal residue of the buckled protomer and *vice versa*, **Fig. 1D**). The communication extends to regions in the CTDs of both protomers and the NTD of the cognate protomer, through localized hotspots at the dimerization interface (**Fig. 1D)**.

In summary, the TRAP1 cysteine which undergoes *S*-nitrosylation, shows a propensity to promote conformational changes to distal sites in the NTD, and at the interface between the two protomers, i.e., in the MD and CTD. Our analysis suggests that *S*-nitrosylation of C516 (human C501) might modulate the activity of the chaperone and not only the propensity of the protein to be degraded by the proteasome, as we previously reported [22].

### 3.2 S-nitrosylation alter protein flexibility and inhibit the functional movements of the chaperone

In light of the PSN results, we wonder if we could identify changes in the dynamics of the TRAP1_SH_ and TRAP1_SNO_ variants that might be related to the conformational changes that the chaperone needs to undergo during the catalytic cycle. As a simple model to address these points, we used MD simulations of the individual protomers for the two TRAP1 variants. This allowed us to see possible motions of opening and closing of the different domains in short simulations of few hundred ns. We compared the simulations using a dimensionality reduction technique based on Principal Component Analysis, highlighting, in such a way, large amplitude motions in the protein [38]. We used the 2D projections of the MD structures along the first and second principal components (**Fig. 2A**). Each point in this *essential* subspace represents a structure of the original MD trajectories. We notice that the *S*-nitrosylation of C516 induced a stiffness in the structure, reducing opening/closing motions and the exploration of a smaller portion of the 2D subspace (**Fig. 2B**). This behavior is especially evident in the TRAP1_SNO_ buckled protomer where the conformational transition between the two states observed in the corresponding TRAP1_SH_ protomer is impaired (**Fig. 2C**).

**Figure 2.**
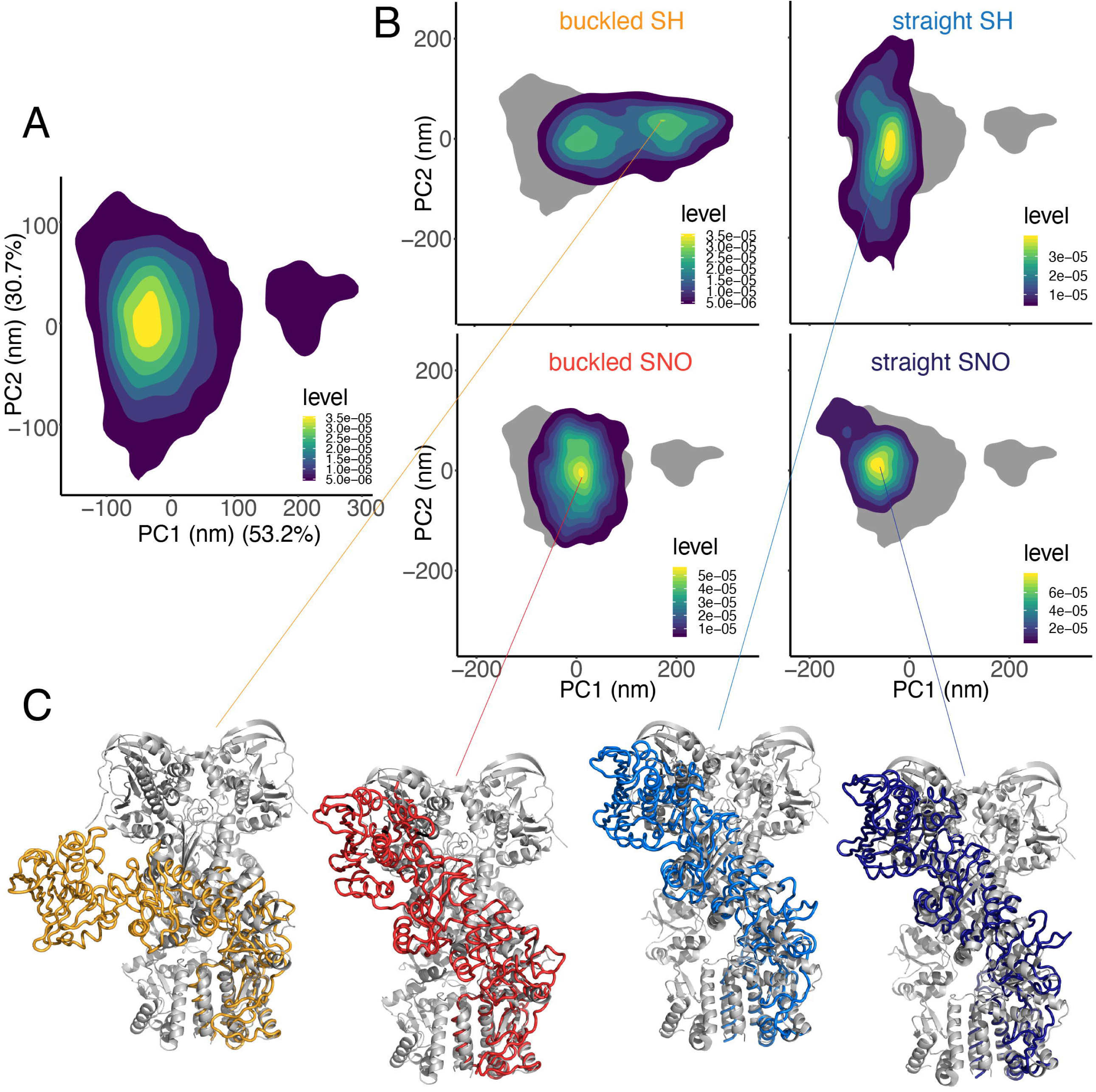
TRAP1 *S*-nitrosylation results in structural stiffness and impairs opening/closing motions, especially in the buckled protomer. **A**) Two-dimensional (2D) density plot, using as reaction coordinates, the first and the second principal components from the Principal Component Analysis (PCA) performed on the Cα atoms. We calculated the PCA on a concatenated trajectory including all the MD simulations of TRAP1_SH_ and TRAP1_SNO_ protomers and performed with GROMOS 54a7 force field. The first two principal components explain more than 83.9 % of the total variance of the system. We report the variance explained by each principal component within the round brackets in the axes of the 2D plot. **B**) Conformational space explored by each MD simulation of TRAP1_SH_ and TRAP1_SNO_ protomers, projected in the essential subspace derived by the PCA. **C**) Conformations collected to identify the most densely populated area in the 2D density plots. Cartoon represent four representative structures of the TRAP1_SH_ and TRAP1_SNO_ buckled and straight protomer (colored in orange, red, light blue and blue, respectively). The structure of the protomers are aligned using as a reference the CTD in the X-ray structure of TRAP1 dimer (colored in grey, PDB entry: 4IPE).

### 3.3 *S*-nitrosylation at Cys501 affects TRAP1 ATPase activity

Based on results so far obtained, we moved to *in vitro* systems, *i.e.* recombinant human TRAP1, to verify *in silico* predictions about the effects induced by *S*-nitrosylation on TRAP1 activity. To this end, we measured ATPase activity by Malachite green phosphate assay upon 4 h incubation with the NO donor SNAP. Colorimetric analyses indicated that SNAP treatment decreased TRAP1 ATPase activity by about 25-30% (**Fig. 3A**). Next, to avoid that the observed phenomenon was related to other possible *S*-nitrosylation-independent reactions induced by NO (e.g. nitration), after SNAP treatment, we dialyzed TRAP1 and incubated it for 1 h with 20 mM ascorbate, a specific SNO reductant. Results shown in **Fig. 3B** indicated that ascorbate addition rescued the activity of the protein, confirming that *S*-nitrosylation was the driving event affecting TRAP1 activity. On the basis of these results, we decided to substitute the nitrosylable Cys (conserved in human TRAP1 at position 501) with a Ser, in order to maintain steric hindrance and polarity. Before mutagenesis, we predicted the effects on protein stability induced by Cys-to-Ser substitution in the form of ΔΔGs. We used the different available crystallographic structures of human and *D. rerio* TRAP1 variants for the calculation and the FoldX energy function (**Table 2**), as we and others showed that this biophysical measurement correlates well with the cellular protein expression levels of protein mutant variants [40–42]. The advantage of using multiple structures of the same protein for this calculation is to overcome the inherent limitation of the FoldX method, which performs only local conformational sampling around the input structure. We noticed that in all the cases, the C516S – and, by homology, C501S – mutation is not predicted to change protein stability, with low ΔΔG values always below the threshold of 1.6 kcal/mol. These calculations suggest that we could use the C501S mutant variant for experimental studies without risking to affect protein expression levels. This conclusion was confirmed by the amount of TRAP1^C501S^ obtained upon purification, which did not differ significantly from TRAP1^WT^ (**Fig. 4A**). TRAP1 variants were than analyzed for their capability to hydrolyze ATP. Malachite green assays showed that TRAP1^C501S^ exhibited a higher ATPase activity than TRAP1^WT^ (**Fig. 4B**) but, most importantly, this was not affected by SNAP addition, confirming that Cys501 nitrosylation affects TRAP1 activity.

**Table 2.** Prediction of changes in protein structural stability upon mutations of C516/C501. The average ΔΔG values and standard deviations associated with protein stability are reported for each TRAP1 crystallographic structure. C516 and C501 refer to the Danio rerio and human TRAP1 variants, respectively. For dimeric structures, the results for each of the monomers (i.e., chain A or B in the PDB file) are reported.

| MUTATION | PDB | $\Delta\Delta G$ (kcal/mol) |
| --- | --- | --- |
| C516S | 4IPE, Chain A | 1 (0.077) |
| C516S | 4IPE, Chain B | 1.62(0.0056) |
| C501S | 4Z1F | 1(0.095) |
| C501S | 4Z1G | 0.56 (0.099) |
| C501S | 4Z1H | -1.41 (0.10) |
| C501S | 5HPH, Chain A | 0.60 (0.00001) |
| C501S | 5HPH, Chain B | 0.85 (0.097) |

**Figure 3.**
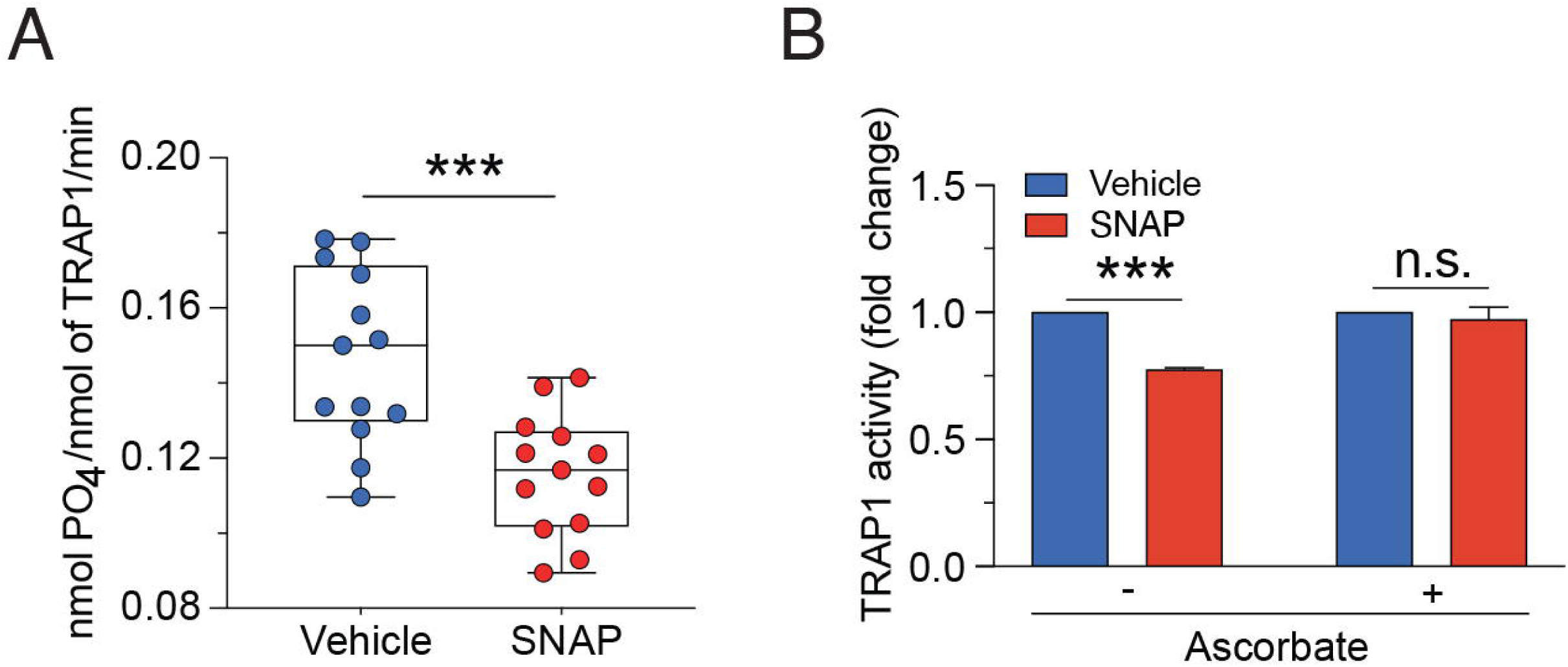
Human recombinant TRAP1 *S*-nitrosylation results in loss of ATPase activity of. **A**) ATPase activity of recombinant TRAP1 was measured as PO_4_^3-^ release by Malachite Green Phosphate Assay following the absorbance at 650 nm in the presence (SNAP) or absence (Vehicle) of the NO donor *S*-nitroso-*N*-acetilpenicilamine (SNAP, 500 μM, 4 h in the dark). Data are shown as nmoles of PO_4_^3-^ released/nmol of TRAP1/min and represent the mean ± SEM of n = 5 independent experiments done in triplicate. \*\*\**p* < 0.001. **B**) ATPase activity as in (A) was conducted in the presence or absence of the SNO-reductant ascorbate. Data are shown as fold change (respect to untreated cells (Vehicle) and represent the mean ± SEM of n = 3 independent experiments done in triplicate. \*\*\**p* < 0.001; n.s., not significant.

**Figure 4.**
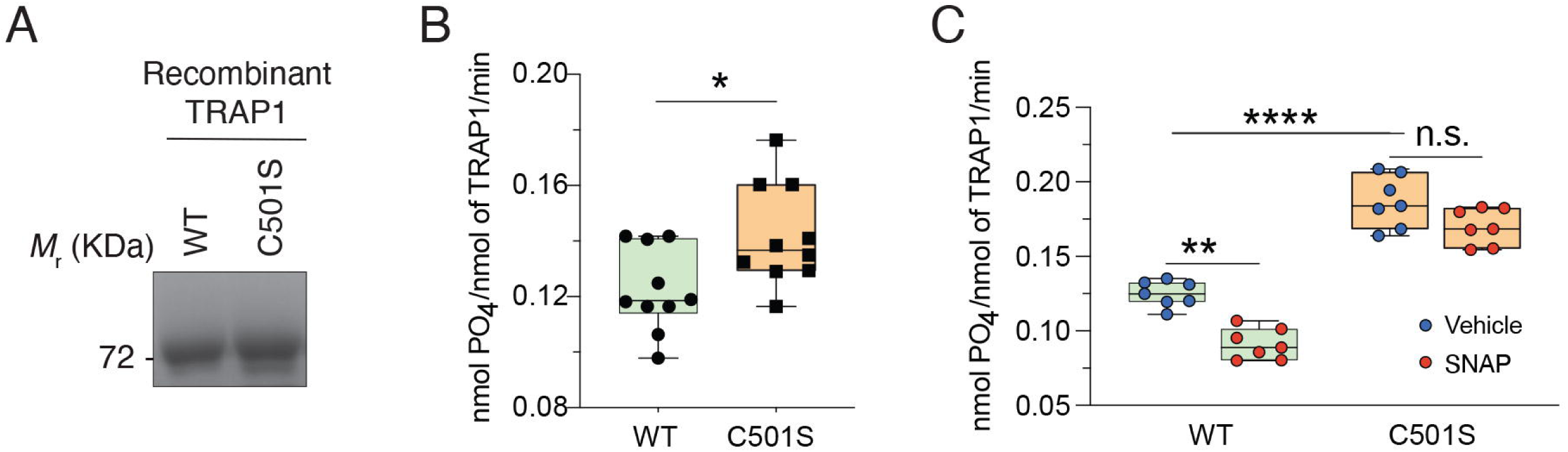
TRAP1 C501S mutant is insensitive to NO-mediated inhibition of ATPase activity. **A**) Coomassie staining of an SDS-PAGE was done to verify purity and yield of WT and C501S mutant form of human recombinant TRAP1. **B**) ATPase activity of recombinant WT and C501S mutant of TRAP1 was measured as PO_4_^3-^ release by Malachite Green Phosphate Assay following the absorbance at 650 nm. Data are shown as nmoles of PO_4_^3-^ released/nmol of TRAP1/min and represent the mean ± SEM of n = 4 independent experiments done in triplicate. \**p* < 0.05. **C**) ATPase activity as in (B) was conducted in the presence (SNAP) or absence (Vehicle) of the NO donor *S*-nitroso-*N*-acetilpenicilamine (SNAP, 500 μM, 4 h in the dark). Data are shown as nmoles of PO_4_^3-^ released/nmol of TRAP1/min and represent the mean ± SEM of n = 4 independent experiments done in triplicate. \*\**p* < 0.01; \*\*\*\**p* < 0.0001; n.s., not significant.

### 3.4 TRAP1 *S*-nitrosylation at Cys501 modulates staurosporin-induced cell death

In order to understand the effects of Cys501 nitrosylation in a biological context, we overexpressed TRAP1^WT^ and TRAP1^C501S^ in HeLa cells (**Fig. 5A**) and analyzed whether the expression of TRAP1^C501S^ could modulate apoptotic response, as this has been reported as one of the processes regulated by TRAP1 [3,15,16,43]. As inducer, we selected staurosporine (STS), as it has been already shown it activates the intrinsic (mitochondrial) pathway of apoptosis [44,45]. Direct cell count upon Trypan blue exclusion indicated that TRAP1^C501S^-expressing cells were more resistant than those carrying the TRAP1^WT^ variant (**Fig. 5B**). This difference was maintained (even increased) upon treatment with NO donors (i.e., DETA-NONOate), that we carried out to induce inhibition TRAP1^WT^ *via S*-nitrosylation. This result was also confirmed by Western blot analyses of caspase 3 and PARP1 cleavage (**Fig. 5C**), which are well-established markers of apoptosis, indicating that the non-nitrosylable form of TRAP1 was protective under these experimental conditions.

**Figure 5.**
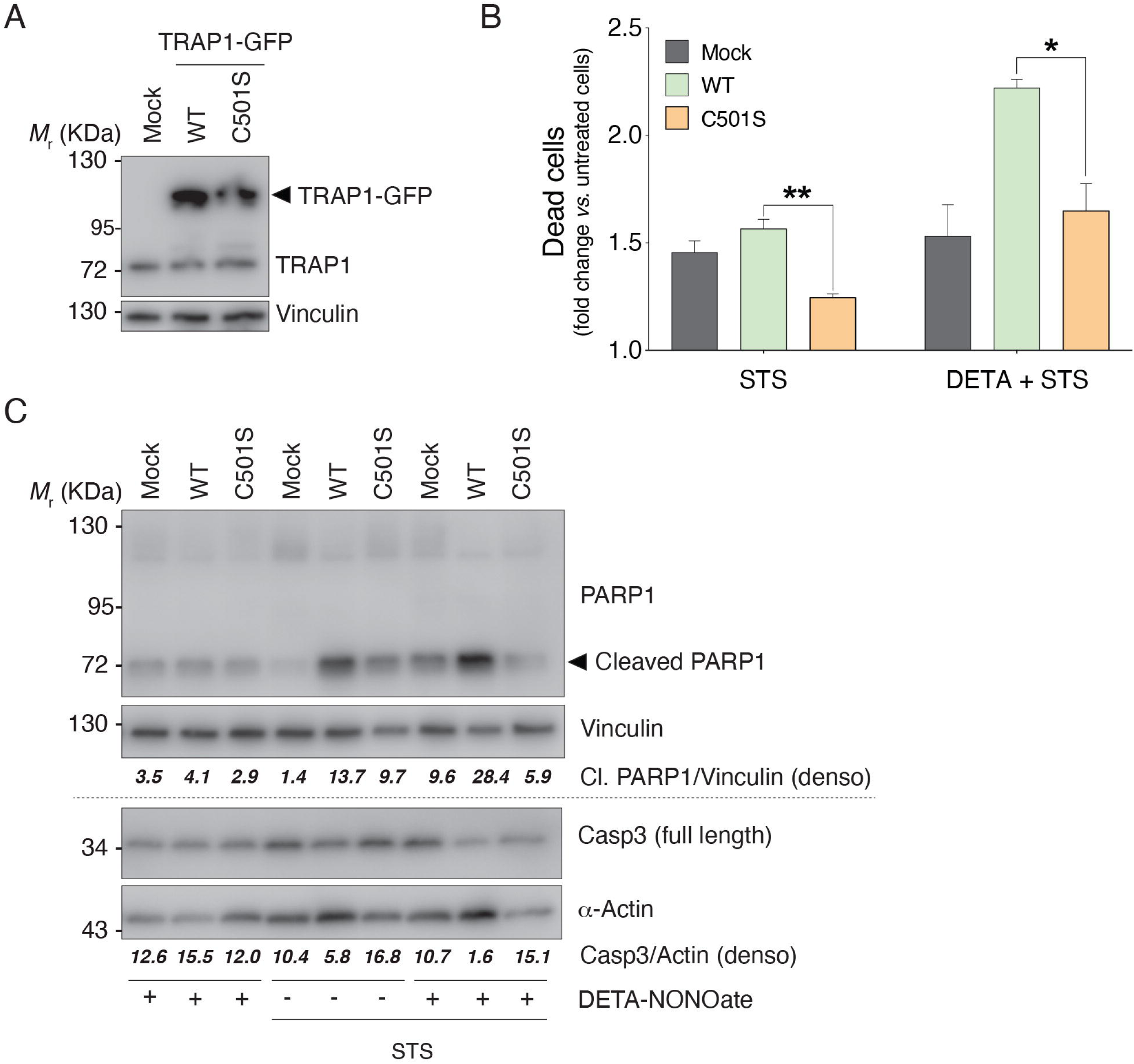
TRAP1 C501S mutant exerts protective role towards staurosporine-induced cell death. **A**) Western blot analyses of HeLa cell lysates were performed to check the expression levels of WT and C501S mutant forms of GFP-fused TRAP1 (TRAP1-GFP) which can be discriminated by the endogenous variant (TRAP1) by the use of an anti-TRAP1 antibody. Western blot shown is representative of at least n = 3 independent experiments giving similar results. **B**) The number of dead cells was evaluated by Trypan blue exclusion test in control cells transfected with an empty vector (Mock), as well as in cell expressing TRAP1^WT^ (WT) and TRAP1^C501S^ (C501S) treated for 4 h with 200 nM staurosporine (STS), in the presence or absence of the NO donor DETA-NONOate (250 μM, maintained for 22-24 h before STS treatment). Data are shown as fold change of dead cells with respect to untreated cells, and represent the mean ± SEM of n =5 independent experiments done in duplicate. \**p* = 0.05; \*\**p* = 0.01. **C**) Western blot analyses of HeLa cells treated as in (B) were performed to evaluate cleaved PARP1 and pro-caspase 3 (Casp3) protein levels. Vinculin and α-actin were used as loading controls. Densitometric analyses (denso) of PARP1/Vinculin or Casp3/actin are shown below each lane. Western blots shown are representative of at least n = 3 independent experiments giving similar results.

## 4. CONCLUSIONS

*S*-nitrosylation of Cys501 is a redox modification associated with TRAP1 degradation *via* proteasome [22]. However, no molecular mechanisms, neither possible structural changes or biological relevance have been provided being related to this, if not the evidence that TRAP1 degradation indirectly causes an increased in SDH expression/activity [22]. In this study we used molecular dynamics simulations and methods based on graph theory and dimensionality reduction to investigate the effects induced by *S-*nitrosylation of Cys501. Our results suggest that the cysteine has an intrinsic propensity to transmit conformational changes to distal sites that are important for the conformational changes of the chaperone during the catalytic cycle or for interaction with the client proteins. Its posttranslational modification is thus likely to perturb the functional dynamics of TRAP1, as we showed using principal component analyses of the simulations. The buckled protomer of the asymmetric TRAP1 structure seems the most sensitive to the modification in our model system and *S*-nitrosylation causes a stiffness of the structure and impairment of motions. This restrains the elements in the C-terminal and middle domain that can work as hinges for these conformational changes. Our data are in good agreement with recent findings from another computational study where the region surrounding the SNO site has been shown with a pronounced sensor signature by perturbation scanning analyses [46], making the area a suitable candidate for allosteric communication. In another computational study using all-atom MD simulations, the authors identified a cross-talk and coupling between the nucleotide site and distal client binding sites, also in the proximity of the *S*-nitrosylation site [23].

The inhibitory effects of Cys501 nitrosylation on ATPase activity support the *in vivo* evidence arguing for a protective role of TRAP1^C501S^ to apoptotic stimuli, mostly if these are associated with NO fluxes. Our data, together with those previously published [22], reinforce the role of Cys501 in maintaining the stability and ensuring the activity of TRAP1. This is in agreement with the higher TRAP1 activity of the mutant, but moves the question on: “why the mutant should be more active?” Work is still ongoing in our laboratory to exactly clarify how *S*-nitrosylation interferes with TRAP1 activity and functions. However, it is reasonable to speculate that, as observed in this work, TRAP1_SNO_ cannot ever reach the “stiff” conformation and, in turn, ever decrease the chaperone efficiency in a population of TRAP1 molecules.

Evolutionarily, TRAP1 is considered the ancestral member of HSP90 family [1] and, for this reason, its activity could probably be more responsive to ROS and NO as hypothesized for many ancestral proteins (i.e., the bacterial transcription factor OxyR) [47,48]. Therefore, this new regulation of TRAP1 could find a rationale if seen from this angle. Whether Cys501 can also act as general “redox antenna” (e.g. whether it can react with H_2_O_2_ and other electrophiles) is still far from being elucidated. However, the evidence that Cys501 is placed in close proximity to another cysteine (Cys527) might suggest the presence of a peroxiredoxin-like intra-molecular redox switch able to function as novel signaling mechanism. Work is in progress in our lab to elucidate this issue and others dealing with the role of S-nitrosylation and redox mechanism underlying TRAP1 regulation. In this regard, preliminary studies suggest that *S*-nitrosylation is a posttranslational modification broadly modulating TRAP1 functions. In particular, mass spectrometry analyses, performed in human TRAP1, indicate that Cys573 (which is not present in TRAP1 from *D. rerio*) can also undergo *S*-nitrosylation, with this event being probably implicated in regulating TRAP1 activity and structure by further, still unknown, mechanisms.

## ACKNOWLEDGEMENTS

We would like to thank Giorgio Colombo’s group to provide the structure of the TRAP1 dimer to use in our simulations and Laila Fisher for secretary assistance. This work has been supported by Danish Cancer Society Grant Kræftens Bekæmpelses Videnskabelige Udvalg (KBVU) Grant [R146-A9414] (to G.F. and E.P.); Associazione Italiana per la Ricerca sul Cancro (AIRC) Grant [IG20719] (to G.F.); the DECI-PRACE 13^th^ HPC Grant CHAPREDO for computational time on Archer (to E.P.); and a LISA CINECA HPC Grant for computational time on Marconi and Pico (HPL13PEJU2, to E.P.); AIRC IG 2017-20749 (to A.R.) and “The neurofibromatosis therapeutic acceleration program” (to A.R.); the Danish National Research Foundation (DNRF125) as fund to the Center of Excellence in Autophagy, Recycling and Disease (CARD).

## Author contributions

F.F., C.P., P.G., and G.C. performed the experiments *in vitro* and cellular experiments, and interpreted the data; F.F., S.R., A.R., G.F., E.P., conceived and designed the study, M.L. J.S.V., M.F.A. and E.P. performed molecular dynamics and *in silico* studies, and interpreted the data; F.P. and A.B. produced the recombinant proteins; F.F., M.L., S.R., J.V., A.B., A.R., E.P. and G.F., critically commented the manuscript and conceived the figures; E.P. and G.F. wrote the manuscript.

## Conflicts of Interest

The authors state that no conflict of interests does exist.

## Notes

https://github.com/ELELAB/TRAP1_activity

https://osf.io/wytg2/

## References

[1] Ho Yeong Song, J.D. Dunbar, Yuan Xin Zhang, D. Guo, D.B. Donner, Identification of a protein with homology to hsp90 that binds the type 1 tumor necrosis factor receptor, J. Biol. Chem. 270 (1995) 3574–3581. doi:10.1074/jbc.270.8.3574.

[2] A. Leskovar, H. Wegele, N.D. Werbeck, J. Buchner, J. Reinstein, The ATPase Cycle of the Mitochondrial Hsp90 Analog Trap1 * □, J. Biol. Chem. 283 (2008) 11677–11688. doi:10.1074/jbc.M709516200.

[3] B.H. Kang, TRAP1 regulation of mitochondrial life or death decision in cancer cells and mitochondria-targeted TRAP1 inhibitors, BMB Rep. 45 (2012) 1–6. doi:10.5483/BMBRep.2012.45.1.1.

[4] A. Joshi, J. Dai, J. Lee, N.M. Ghahhari, G. Segala, K. Beebe, F.T.F. Tsai, L. Neckers, D. Picard, The mitochondrial HSP90 paralog TRAP1 forms an OXPHOS-regulated tetramer and is involved in maintaining mitochondrial metabolic homeostasis, BioRxiv. (2019) 679431. doi:10.1101/679431.

[5] L.A. Lavery, J.R. Partridge, T.A. Ramelot, D. Elnatan, M.A. Kennedy, D.A. Agard, Structural Asymmetry in the Closed State of Mitochondrial Hsp90 (TRAP1) Supports a Two-Step ATP Hydrolysis Mechanism, Mol. Cell. 53 (2014) 330–343. doi:10.1016/j.molcel.2013.12.023.

[6] I. Masgras, C. Sanchez-Martin, G. Colombo, A. Rasola, The Chaperone TRAP1 As a Modulator of the Mitochondrial Adaptations in Cancer Cells, Front. Oncol. 7 (2017) 1–10. doi:10.3389/fonc.2017.00058.

[7] J.R. Partridge, L.A. Lavery, D. Elnatan, N. Naber, R. Cooke, D.A. Agard, A novel N-terminal extension in mitochondrial TRAP1 serves as a thermal regulator of chaperone activity, Elife. (2014) 1–21. doi:10.7554/eLife.03487.

[8] L.A. Lavery, J.R. Partridge, T.A. Ramelot, D. Elnatan, M.A. Kennedy, D.A. Agard, Article Structural Asymmetry in the Closed State a Two-Step ATP Hydrolysis Mechanism, Mol. Cell. 53 (2014) 330–343. doi:10.1016/j.molcel.2013.12.023.

[9] N. Sung, J. Lee, J.-H. Kim, C. Chang, A. Joachimiak, S. Lee, F.T.F. Tsai, Mitochondrial Hsp90 is a ligand-activated molecular chaperone coupling ATP binding to dimer closure through a coiled-coil intermediate, Proc Natl Acad Sci U S A. 113 (2016) 2952–2957. doi:10.1073/pnas.1516167113.

[10] D. Elnatan, M. Betegon, Y. Liu, T. Ramelot, M.A. Kennedy, D.A. Agard, Symmetry broken and rebroken during the ATP hydrolysis cycle of the mitochondrial Hsp90 TRAP1, Elife. 6 (2017) e25235. doi:10.7554/eLife.25235.

[11] S. Yoshida, S. Tsutsumi, G. Muhlebach, C. Sourbier, M.-J. Lee, S. Lee, E. Vartholomaiou, M. Tatokoro, K. Beebe, N. Miyajima, R.P. Mohney, Y. Chen, H. Hasumi, W. Xu, H. Fukushima, K. Nakamura, F. Koga, K. Kihara, J. Trepel, D. Picard, L. Neckers, Molecular chaperone TRAP1 regulates a metabolic switch between mitochondrial respiration and aerobic glycolysis., Proc. Natl. Acad. Sci. U. S. A. 110 (2013) E1604–12. doi:10.1073/pnas.1220659110.

[12] M.R. Amoroso, D.S. Matassa, L. Sisinni, G. Lettini, M. Landriscina, F. Esposito, TRAP1 revisited: Novel localizations and functions of a “next-generation” biomarker (Review), Int. J. Oncol. 45 (2014) 969–977. doi:10.3892/ijo.2014.2530.

[13] A. Rasola, L. Neckers, D. Picard, Mitochondrial oxidative phosphorylation TRAP(1)ped in tumor cells, Trends Cell Biol. 24 (2014) 455–463. doi:10.1016/j.tcb.2014.03.005.

[14] Y.C. Chae, M.C. Caino, S. Lisanti, J.C. Ghosh, T. Dohi, N.N. Danial, J. Villanueva, S. Ferrero, V. Vaira, L. Santambrogio, S. Bosari, L.R. Languino, M. Herlyn, D.C. Altieri, Control of Tumor Bioenergetics and Survival Stress Signaling by Mitochondrial HSP90s, Cancer Cell. 22 (2012) 331–344. doi:10.1016/j.ccr.2012.07.015.

[15] E. Costantino, F. Maddalena, S. Calise, A. Piscazzi, V. Tirino, A. Fersini, A. Ambrosi, V. Neri, F. Esposito, M. Landriscina, TRAP1, a novel mitochondrial chaperone responsible for multi-drug resistance and protection from apoptotis in human colorectal carcinoma cells, Cancer Lett. 279 (2009) 39–46. doi:10.1016/j.canlet.2009.01.018.

[16] B.H. Kang, J. Plescia, T. Dohi, J. Rosa, S.J. Doxsey, D.C. Altieri, Regulation of Tumor Cell Mitochondrial Homeostasis by an Organelle-Specific Hsp90 Chaperone Network, Cell. 131 (2007) 257–270. doi:10.1016/j.cell.2007.08.028.

[17] M. Sciacovelli, G. Guzzo, V. Morello, C. Frezza, L. Zheng, N. Nannini, F. Calabrese, G. Laudiero, F. Esposito, M. Landriscina, P. Defilippi, P. Bernardi, A. Rasola, The mitochondrial chaperone TRAP1 promotes neoplastic growth by inhibiting succinate dehydrogenase, Cell Metab. 17 (2013) 988–999. doi:10.1016/j.cmet.2013.04.019.

[18] M.A. Selak, S.M. Armour, E.D. MacKenzie, H. Boulahbel, D.G. Watson, K.D. Mansfield, Y. Pan, M.C. Simon, C.B. Thompson, E. Gottlieb, Succinate links TCA cycle dysfunction to oncogenesis by inhibiting HIF-α prolyl hydroxylase, Cancer Cell. 7 (2005) 77–85. doi:10.1016/j.ccr.2004.11.022.

[19] J.W. Pridgeon, J.A. Olzmann, L.S. Chin, L. Li, PINK1 protects against oxidative stress by phosphorylating mitochondrial chaperone TRAP1, PLoS Biol. 5 (2007) 1494–1503. doi:10.1371/journal.pbio.0050172.

[20] I. Masgras, F. Ciscato, A.M. Brunati, E. Tibaldi, S. Indraccolo, M. Curtarello, F. Chiara, G. Cannino, E. Papaleo, M. Lambrughi, G. Guzzo, A. Gambalunga, M. Pizzi, V. Guzzardo, M. Rugge, S.E. Vuljan, F. Calabrese, P. Bernardi, A. Rasola, Absence of Neurofibromin Induces an Oncogenic Metabolic Switch via Mitochondrial ERK-Mediated Phosphorylation of the Chaperone TRAP1, Cell Rep. 18 (2017) 659–672. doi:10.1016/j.celrep.2016.12.056.

[21] H.-K. Park, J.-H. Hong, Y.T. Oh, S.S. Kim, J. Yin, A.-J. Lee, Y.C. Chae, J.H. Kim, S.-H. Park, C.-K. Park, M.-J. Park, J.B. Park, B.H. Kang, Interplay between TRAP1 and Sirtuin-3 Modulates Mitochondrial Respiration and Oxidative Stress to Maintain Stemness of Glioma Stem Cells, Cancer Res. 79 (2019) 1369 LP – 1382. doi:10.1158/0008-5472.CAN-18-2558.

[22] S. Rizza, C. Montagna, S. Cardaci, E. Maiani, G. Di Giacomo, V. Sanchez-Quiles, B. Blagoev, A. Rasola, D. De Zio, J.S. Stamler, F. Cecconi, G. Filomeni, S-nitrosylation of the mitochondrial chaperone TRAP1 sensitizes hepatocellular carcinoma cells to inhibitors of succinate dehydrogenase, Cancer Res. 76 (2016) 4170–4182. doi:10.1158/0008-5472.CAN-15-2637.

[23] E. Moroni, D.A. Agard, G. Colombo, The Structural Asymmetry of Mitochondrial Hsp90 (Trap1) Determines Fine Tuning of Functional Dynamics, J. Chem. Theory Comput. 14 (2018) 1033–1044. doi:10.1021/acs.jctc.7b00766.

[24] B. Webb, A. Sali, Comparative Protein Structure Modeling Using MODELLER, Curr. Protoc. Bioinforma. 54 (2016) 5.6.1-5.6.37. doi:10.1002/cpbi.3.

[25] E.Z. Eisenmesser, O. Millet, W. Labeikovsky, D.M. Korzhnev, M. Wolf-Watz, D.A. Bosco, J.J. Skalicky, L.E. Kay, D. Kern, Intrinsic dynamics of an enzyme underlies catalysis, Nature. 438 (2005) 117–121. doi:10.1038/nature04105.

[26] B. Ma, R. Nussinov, Conformational footprints, Nat. Chem. Biol. 12 (2016) 890–891. doi:10.1038/nchembio.2212.

[27] C. Margreitter, D. Petrov, B. Zagrovic, Vienna-PTM web server: a toolkit for MD simulations of protein post-translational modifications, Nucleic Acids Res. 41 (2013) W422–W426. doi:10.1093/nar/gkt416.

[28] B. Hess, C. Kutzner, D. van der Spoel, E. Lindahl, GROMACS 4:□ Algorithms for Highly Efficient, Load-Balanced, and Scalable Molecular Simulation, J. Chem. Theory Comput. 4 (2008) 435–447. doi:10.1021/ct700301q.

[29] P. Robustelli, S. Piana, D.E. Shaw, Developing a molecular dynamics force field for both folded and disordered protein states, Proc. Natl. Acad. Sci. 115 (2018) E4758 LP–E4766. doi:10.1073/pnas.1800690115.

[30] N. Schmid, A.P. Eichenberger, A. Choutko, S. Riniker, M. Winger, A.E. Mark, W.F. van Gunsteren, Definition and testing of the GROMOS force-field versions 54A7 and 54B7, Eur. Biophys. J. 40 (2011) 843. doi:10.1007/s00249-011-0700-9.

[31] D. Petrov, C. Margreitter, M. Grandits, C. Oostenbrink, B. Zagrovic, A Systematic Framework for Molecular Dynamics Simulations of Protein Post-Translational Modifications, PLOS Comput. Biol. 9 (2013) e1003154. doi:10.1371/journal.pcbi.1003154.

[32] S. Han, Force field parameters for S-nitrosocysteine and molecular dynamics simulations of S-nitrosated thioredoxin, Biochem. Biophys. Res. Commun. 377 (2008) 612–616. doi:10.1016/j.bbrc.2008.10.017.

[33] T.J. Dolinsky, P. Czodrowski, H. Li, J.E. Nielsen, J.H. Jensen, G. Klebe, N.A. Baker, PDB2PQR: expanding and upgrading automated preparation of biomolecular structures for molecular simulations, Nucleic Acids Res. 35 (2007) W522–W525. doi:10.1093/nar/gkm276.

[34] M. Tiberti, G. Invernizzi, M. Lambrughi, Y. Inbar, G. Schreiber, E. Papaleo, PyInteraph: A Framework for the Analysis of Interaction Networks in Structural Ensembles of Proteins, J. Chem. Inf. Model. 54 (2014) 1537–1551. doi:10.1021/ci400639r.

[35] J. Salamanca Viloria, M.F. Allega, M. Lambrughi, E. Papaleo, An optimal distance cutoff for contact-based Protein Structure Networks using side-chain centers of mass, Sci. Rep. 7 (2017) 2838. doi:10.1038/s41598-017-01498-6.

[36] E. Papaleo, Integrating atomistic molecular dynamics simulations, experiments, and network analysis to study protein dynamics: strength in unity, Front. Mol. Biosci. 2 (2015) 28. doi:10.3389/fmolb.2015.00028.

[37] R. Guerois, J.E. Nielsen, L. Serrano, Predicting Changes in the Stability of Proteins and Protein Complexes: A Study of More Than 1000 Mutations, J. Mol. Biol. 320 (2002) 369–387. doi:10.1016/S0022-2836(02)00442-4.

[38] A. Amadei, A.B.M. Linssen, H.J.C. Berendsen, Essential dynamics of proteins, Proteins Struct. Funct. Bioinforma. 17 (1993) 412–425. doi:10.1002/prot.340170408.

[39] O.H. Lowry, N.J. Rosebrough, A.L. Farr, R.J. Randall, PROTEIN MEASUREMENT WITH THE FOLIN PHENOL REAGENT, J. Biol. Chem.. 193 (1951) 265–275. http://www.jbc.org/content/193/1/265.short.

[40] A.B. Abildgaard, A. Stein, S. V Nielsen, K. Schultz-Knudsen, E. Papaleo, A. Shrikhande, E.R. Hoffmann, I. Bernstein, A.-M. Gerdes, M. Takahashi, C. Ishioka, K. Lindorff-Larsen, R. Hartmann-Petersen, Computational and cellular studies reveal structural destabilization and degradation of MLH1 variants in Lynch syndrome, Elife. 8 (2019) e49138. doi:10.7554/eLife.49138.

[41] R. Scheller, A. Stein, S. V Nielsen, F.I. Marin, A.-M. Gerdes, M. Di Marco, E. Papaleo, K. Lindorff-Larsen, R. Hartmann-Petersen, Toward mechanistic models for genotype–phenotype correlations in phenylketonuria using protein stability calculations, Hum. Mutat. 40 (2019) 444–457. doi:10.1002/humu.23707.

[42] S. V Nielsen, A. Stein, A.B. Dinitzen, E. Papaleo, M.H. Tatham, E.G. Poulsen, M.M. Kassem, L.J. Rasmussen, K. Lindorff-Larsen, R. Hartmann-Petersen, Predicting the impact of Lynch syndrome-causing missense mutations from structural calculations, PLOS Genet. 13 (2017) e1006739. doi:10.1371/journal.pgen.1006739.

[43] M. Landriscina, G. Laudiero, F. Maddalena, M.R. Amoroso, A. Piscazzi, F. Cozzolino, M. Monti, C. Garbi, A. Fersini, P. Pucci, F. Esposito, Mitochondrial Chaperone Trap1 and the Calcium Binding Protein Sorcin Interact and Protect Cells against Apoptosis Induced by Antiblastic Agents, Cancer Res. 70 (2010) 6577 LP – 6586. doi:10.1158/0008-5472.CAN-10-1256.

[44] M. Malsy, D. Bitzinger, B. Graf, A. Bundscherer, Staurosporine induces apoptosis in pancreatic carcinoma cells PaTu 8988t and Panc-1 via the intrinsic signaling pathway, Eur. J. Med. Res. 24 (2019) 5. doi:10.1186/s40001-019-0365-x.

[45] J. Manns, M. Daubrawa, S. Driessen, F. Paasch, N. Hoffmann, A. Löffler, K. Lauber, A. Dieterle, S. Alers, T. Iftner, K. Schulze-Osthoff, B. Stork, S. Wesselborg, Triggering of a novel intrinsic apoptosis pathway by the kinase inhibitor staurosporine: activation of caspase-9 in the absence of Apaf-1, FASEB J. 25 (2011) 3250–3261. doi:10.1096/fj.10-177527.

[46] G.M. Verkhivker, Dynamics-based community analysis and perturbation response scanning of allosteric interaction networks in the TRAP1 chaperone structures dissect molecular linkage between conformational asymmetry and sequential ATP hydrolysis, Biochim. Biophys. Acta - Proteins Proteomics. 1866 (2018) 899–912. doi:10.1016/j.bbapap.2018.04.008.

[47] C.T. Stomberski, D.T. Hess, J.S. Stamler, Protein S-Nitrosylation: Determinants of Specificity and Enzymatic Regulation of S-Nitrosothiol-Based Signaling, Antioxid. Redox Signal. 30 (2019) 1331–1351. doi:10.1089/ars.2017.7403.

[48] D. Seth, A. Hausladen, Y.-J. Wang, J.S. Stamler, Endogenous Protein S-Nitrosylation in E. coli: Regulation by OxyR, Science. 336 (2012) 470 LP – 473. doi:10.1126/science.1215643.

